# On Signalling and Estimation Limits for Molecular Birth-Processes

**DOI:** 10.1101/319889

**Authors:** Kris V Parag

## Abstract

Understanding and uncovering the mechanisms or motifs that molecular networks employ to regulate noise is a key problem in cell biology. As it is often difficult to obtain direct and detailed insight into these mechanisms, many studies instead focus on assessing the best precision attainable on the signalling pathways that compose these networks. Molecules signal one another over such pathways to solve noise regulating estimation and control problems. Quantifying the maximum precision of these solutions delimits what is achievable and allows hypotheses about underlying motifs to be tested without requiring detailed biological knowledge. The pathway capacity, which defines the maximum rate of transmitting information along it, is a widely used proxy for precision. Here it is shown, for estimation problems involving elementary yet biologically relevant birth-process networks, that capacity can be surprisingly misleading. A time-optimal signalling motif, called birth-following, is derived and proven to better the precision expected from the capacity, provided the maximum signalling rate constraint is large and the mean one above a certain threshold. When the maximum constraint is relaxed, perfect estimation is predicted by the capacity. However, the true achievable precision is found highly variable and sensitive to the mean constraint. Since the same capacity can map to different combinations of rate constraints, it can only equivocally measure precision. Deciphering the rate constraints on a signalling pathway may therefore be more important than computing its capacity.

## 1. Introduction

In cell biology small molecular populations signal one another, by modulating the rates at which they react (e.g. via catalysis), to solve regulatory estimation and control problems [1]. These modulations are intrinsically noisy due to fluctuations arising from the random timing of birth (synthesis) and death (turnover) reactions [2]. ‘Signals’ here are stochastic changes in population size that encode information about some target molecule of interest, with population size representing, for example, gene copy numbers or messenger protein counts. Intracellular signalling is therefore, fundamentally, a noisy information theoretic problem [3].

Since biochemical networks attenuate, filter, or utilise this noise to function precisely [4], much research has focussed on demystifying what constitutes effective signalling, and on what sets the limits of cellular precision [5] [6] [7]. ‘Precision’ refers to a mean squared error (mse) or distortion that measures how well a regulatory network solves relevant estimation or control problems, despite intrinsic noise. Higher precision implies smaller mse. Understanding, uncovering and specifying the limits of achievable precision helps delineate what cells can and cannot do, better quantifies the impact of noise, aids the identification of optimal signalling motifs and greatly constrains the assumptions made when validating models against experimental data [8] [5].

Consider a signalling molecule tasked with estimating or controlling some target molecule. A signalling pathway is then the route of information flow from the target to signalling species. The pathway capacity, which defines the maximum rate at which information can be transmitted along it, is believed to uniquely control the best achievable precision [7] [6]. Recent analyses have computed these capacities from empirically derived molecular distributions by modelling the signalling pathway as a Gaussian channel [9] [10] [11]. This assumes that received signals are continuous waveforms corrupted by additive white noise [7] [5]. Channel capacity is then determined by a single parameter, the signal to noise ratio (snr) [12], and has a unique inverse relationship to the best mse achievable [13].

While this approach simplifies analysis and has yielded many insights, the link between information and precision remains unresolved [9]. One reason for this is that Gaussian channels cannot properly describe discrete information structures [14]. In cellular signalling both the information and noise manifest in the timing of discrete events [15]. Noise is also not additive. These issues were addressed in [1], where the Poisson channel was proposed as a replacement. Poisson channels convert a continuous time input into a Poisson point process output with intensity equal to that input [16]. Here the input is the rate of the signalling reaction, which encodes information about the target molecule of interest, and the output is a discrete signal event stream that embeds this information within the timing of its events.

Constraints on this input limit the Poisson channel capacity, and so imply more realistic lower bounds on the achievable mse. These bounds, which were derived in [1], preserve the discreteness of the signalling molecule and significantly advanced the understanding of the information-precision relationship. However, they are not perfect and are still being investigated [8]. In particular, they approximate fluctuations in the target molecule population size with a continuous diffusion process. A heuristic, discrete encoding strategy, known as birth-following, was subsequently found to achieve smaller mse values than these bounds, provided the maximum signalling rate is relatively large and nonlinear strategies are allowed [17] [18]. Under these conditions, target molecule discreteness matters.

This result raises two questions, which form the motivation and subject matter of this work. First, if no continuity approximations are made, what is the optimal signalling encoder that is also biologically feasible? Deriving such an encoder would (i) reveal the best mechanism for embedding signal-timing information, which could be useful for synthetic circuit design [19] and (ii) provide a test-bed for contextualising actually achievable precision against the best known bounds from [1]. This is important as it is not known whether any realisable encoder can actually attain the precision stipulated by the capacity of a signalling pathway [12]. If none can, then existing bounds overestimate how precisely regulatory problems can be solved.

This question is investigated in the context of birth-process estimation, in which both the target and signalling molecules have no turnover, and the signalling population size tries to track or monitor that of the target. This framework models the behaviour of stable, long-lived molecules such as scaffold or timekeeper proteins, and has been used in recent studies such as [15] and [20]. Birth-processes are still not completely understood and form elementary components of many complex regulatory networks. Birth-following, which asserts the maximum signalling rate whenever there are excess target molecules to estimate, will be found to emerge as the optimal, minimum time, signalling encoder for this entire class of problems.

Second, is capacity always a reliable proxy for precision? Or do channel constraints more strongly shape optimal encoding and realisable precision? Poisson channel capacity is a function of both the mean and maximum signalling rate constraints, denoted 〈*f*〉 and *f*_max_ [21] (see Eq. (2)). The practical influence of these constraints has been understudied [22], and is not as intuitive as the Gaussian snr [23]. Over a wide range of estimation problems, it will be shown that birth-following can consistently achieve a mse smaller than the bounds of [1], which supposedly underestimate the minimum achievable mse. This greatly generalises the results in [18] and holds when *f*_max/〈*f*〉_ ≫ 1 and 〈*f*〉 ≥ *u* with *u* as the birth rate of the target species of interest.

Under these conditions the capacity of the Poisson channel and the speed of signalling are sufficiently large that fast jumps in the target cannot be approximated with continuous functions. If *f*_max_ is increased further an unlimited Poisson channel capacity can be attained. In this information rich setting perfect precision would be expected. However, it will be proven that unless 〈*f*〉 ≥ *u* is guaranteed, it is impossible, even with optimal encoders, to obtain perfect precision. This interesting observation holds even when the target is made a diffusion process so that no approximations exist.

Consequently, it is the Poisson channel constraints and not its capacity that sets the limits of precision. In fact a given Poisson capacity can map to many combinations of mean and maximum signalling constraints, leading to equivocal notions of achievable precision. This is in contrast to the bijective relationships between mse, snr and capacity on Gaussian channels [13]. Higher capacities therefore do not necessitate better performance and the relationship between information and precision, when discreteness matters, is not simple, even for the simplest regulatory network.

## 2. Methods

### 2.1. Estimation Problem Framework

The molecular estimation problem is defined here, and its links to signalling and information are established. Let 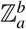 denote the integer set {*a, a* + 1,…, *b* − 1, *b*} with *b* > *a*. The target or estimated molecule is *X*_1_ and has integer population size at time *t* ≥ 0 of *x*_1_(*t*). For convenience, the *t* index will usually be dropped. The causal size history of *x*_1_ over 0 ≤ *s* ≤ *t* is 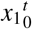. The signalling molecule is *X_j_*, 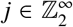, with size *x_j_*(*t*) and history 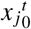. Often *j* = 2, with *j* > 2 only used when complex signalling networks are considered. *X*_1_, however, always represents the target species. Populations fluctuate randomly in time due to Markov birth, *x_i_* → *x_i_* + 1 (denoted 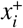) or death, 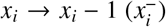, reactions. It is assumed that *X*_1_ cannot be directly observed, and information about it is only available through *X_j_*, which has a birth or signalling rate, *f* that has full knowledge of 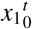 and 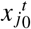 (see Eq. (1) below). Causally estimating *x*_1_ given the observable signalling history 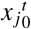 is the main mathematical problem of interest in this work.

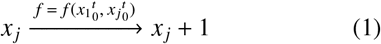

As observed in [1], Eq. (1) also describes a Poisson channel that converts a non-negative input, *f*, into a signal output, *x_j_* [16]. Here *x_j_* is a Poisson process, and *f* can be thought of as the channel encoder responsible for converting *x*_1_ into a form suitable for communication. On this channel information about *x*_1_ is embedded in the timing of *x_j_* birth events, implying a natural link between timing accuracy and the mutual information, denoted 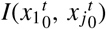. The capacity, 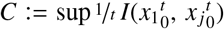, delimits the maximum rate at which information can be transmitted across this channel [1]. Higher capacities allow for more accurate event timing, and the supremum here is taken over all possible 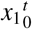 distributions.

The channel decoder, *g*, is some function that reconstructs *x*_1_ from the channel output (observed *x*_2_ event stream). It generates the estimate 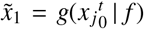. Estimate precision is measured using the mean squared error (mse) distortion, 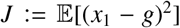, with the minimum mse (mmse) as *J*\*. The mmse is achieved by the optimal decoder 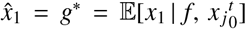 [24]. However, *g*\* is often analytically intractable. As a result, the *x*_1_ estimation problem is considered solved when a good *f* and *g* pair, which leads to a small *J*, is found. Such a pair is called a codec. The explicit dependence of *g* on *f* will usually be dropped for simplicity. Fig. 1 summarises the causal *x*_1_ estimation problem.

**Figure 1:**
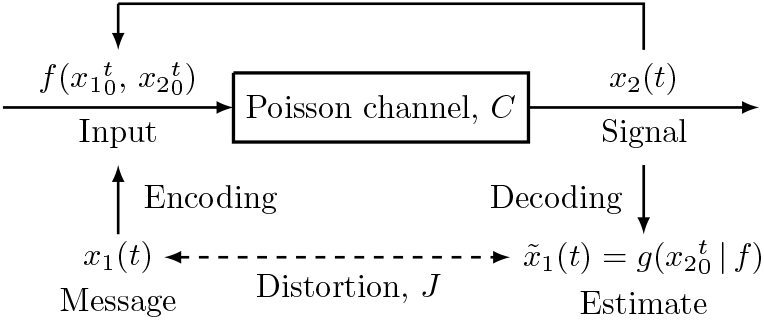
Capacity-distortion schematic. An estimation problem involving two molecular species is specified. The target population, *x*_1_(*t*), is encoded (with feedback) using 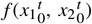 and then sent across a Poisson channel with finite capacity, *C*. This resulting output signal *x*_2_(*t*) is then decoded with 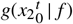 to obtain an estimate 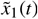. The mean squared error distortion between the original message and the decoded reconstruction is *J*. Higher *C* should imply smaller *J*.

### 2.2. Capacity Based Precision Bounds

The capacity of the channel in Fig. 1 is determined by the signalling constraints placed on *f*. It is common for mean 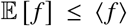 and maximum max(*f*) ≤ *f*_max_ constraints to be considered, leading to Eq. (2) [16].

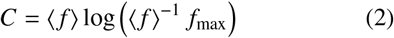

Eq. (2) quantifies the link between signalling (which controls *x_j_* event timing) and information. The mean-maximum properties of the encoder appear fundamental since Eq. (2) holds when the system is generalised so that *x_j_* also modulates *x*_1_ births, and is only scaled by *p*/*p*−1 [21] when the maximum constraint is relaxed to the norm condition 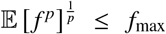 for 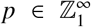. These constraints represent biological limitations for signalling on a pathway, and are usually preset. The sensitivity of precision to both the constraints and the capacity forms the main line of investigation in this work.

A major result of [1], was the realisation that this finite signalling-capacity relationship implies a limit on how small *J*\* can be made. This limit is known as the distortion or capacity bound, *D*, and satisfies *J*\* ≥ *D*. Here *D* is the mse that defines how precisely *X*_1_ could be estimated if it obeyed a diffusion process with equivalent mean (since *X*_1_ is actually Poisson this is known as a diffusion approximation). The relationship between *J*\* and *D* follows from a mutual information (data processing) inequality [12]. Further details of this derivation can be found in the supplement of [1].

If *X*_1_ reacts as 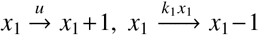 then Eq. (3), with *u* and *k*_1_ as rate constants and *w* a standard Wiener process, is the appropriate diffusion approximation.

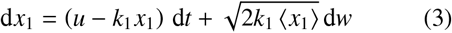

The relevant distortion bound follows from [1] with its dependence on *C* deduced using the information-distortion properties of the diffusion process, under a constraint that *x*_2_ can only depend causally on *x*_1_. This yields Eq. (4) with *J* as the mse for any arbitrary en-coder and *k*_1_ 〈*x*_1_〉 = 〈*u*〉 due to equilibrium.

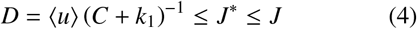

Eq. (4) holds for all non-linear encoders and is a known conservative lower limit on the mmse. Tighter expressions do exist but are only valid for linear encoders, and so not pertinent here [1]. The inverse dependence between *D* and *C* is analogous to the well known Gaussian channel capacity-distortion relationship [13].

In this work Eq. (3) and Eq. (4) will be modified and adapted to derive capacity bounds for various signalling-estimation problems. These *D* functions will then be compared to the achievable *J* or *J*\* for each problem. As in [18] the performance ratio *ψ*:= *j/D* (or *ψ*\* ≤ *ψ* when *J*\* is available) will be used to measure the goodness of a codec. If *ψ* is close to 1 then the codec attaining *J* is seen as good since it is close to the supposed best achievable distortion. Critically, if *ψ*\* < 1 then *J*\* < *D*, which violates the expected relationship. In such cases the capacity actually underestimates the attainable precision on a signalling pathway.

### 2.3. Birth-Following Estimation

Birth-following was introduced in [17] and [18] as a heuristic solution to some of the estimation problems of Section 2.1. The main results from those papers are condensed and extended here, into a key theorem. Consider the simplest non-trivial reaction scheme involving two species, *X*_1_ and *X*_2_, no deaths and 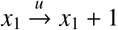 with *u* as a constant. Then *x*_1_ ~ Po(*u*) (Poisson distribution with mean 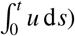. The *X*_2_ signalling reaction is Eq. (1) so *x*_2_ ~ Po(*f*) [25]. This reaction set is an elementary motif for more complex signalling networks. The diffusion approximation for *x*_1_ is given in Eq. (5).

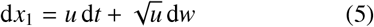

To solve the *x*_1_ estimation problem *f* must be designed, within its mean-maximum constraints. Piecewise continuous processes, which switch between 0 and *f*_max_ achieve Poisson channel capacity [26]. The memoryless birth-following encoder [17], inspired by this, signals the most natural aspect of a birth-process. This encoder only uses the current population size difference *e* = *x*_1_ − *x*_2_, so that 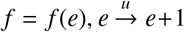 and 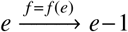. Birth-following is then defined as in Eq. (6) and eventually signals every 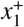 with an 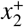, as shown in Fig. 2.

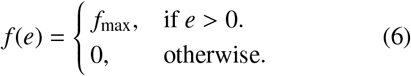

**Figure 2:**
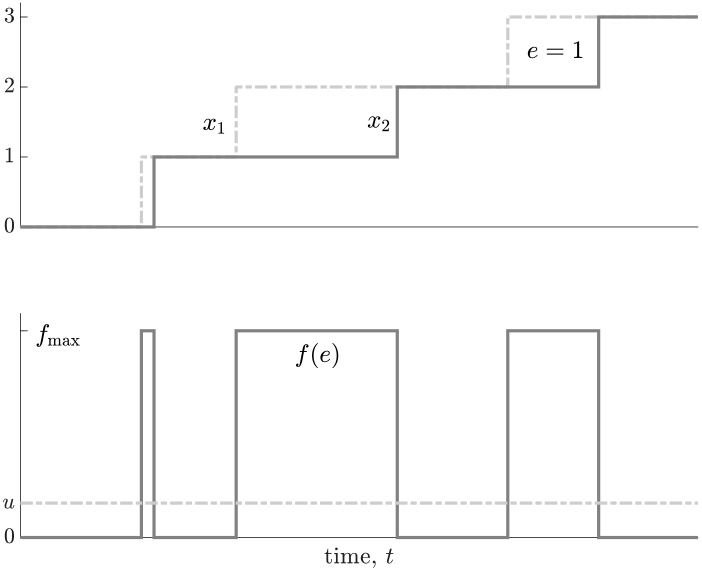
Birth-following encoding. The target molecule, *X*_1_ has synthesis rate, *u* (both dashed). The signalling molecule, *X*_2_, encodes information (about *X*_1_) via its synthesis rate, *f*(*e*) with *e* = *x*_1_ − *x*_2_ (both solid). The top panel shows molecular counts and the bottom one illustrates synthesis rates. Birth-following sets *f*(*e*) to *f*_max_ > *u* whenever there are remaining *X*_1_ molecules to signal (i.e. *e* > 0), else *f* = 0. As *ρ* = *u*/*f*_max_ decreases the gaps between *x*_1_ and *x*_2_ shrink.

An *M*|*M*|1 queue (the ‘M’ stands for Markov, ‘1’ indicates a single server) is a process in which customers randomly arrive with some Poisson rate *u* and are then served with Poisson rate *f*_max_ [27]. The ability of this queue to efficiently serve customers is described by its utilisation, *ρ* = *u*/*f*_max_ < 1. If *e* is interpreted as a number of queueing customers, then reactions 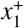 and 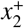 respectively represent arriving and departing customers. Using this analogy, birth-following encoding exactly describes an *M*|*M*|1 queue with *u* = 〈*f*〉. This allows the simplification of Eq. (4) into Eq. (7) below [17].

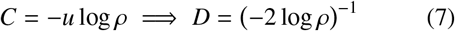

The main theorem can now be stated, in terms of achievable performance, relative to the bound in Eq. (7).

#### Theorem 1.

The birth-following encoder (Eq. (6)) and the memoryless decoder 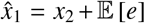 form an asymptotically optimal codec for simple birth-processes, and achieve a relative performance ratio *ψ*\* < 1 when the utilisation *ρ* is small, due to a factor −*ρ* log *ρ*.

Theorem 1 is proven using central results from [17] and [18]. The optimal decoder and its mmse are 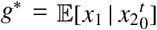 and 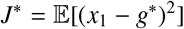 24]. Expanding this and substituting for *e* gives 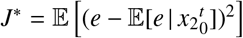. A corollary of Burke’s theorem [28] states that the number of customers in any *M*|*M*|*n* queue (‘*n*’ is the number of servers and *n* = 1 here) is independent of its past departures [28]. This implies that 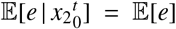.

Consequently, the optimal causal decoder is 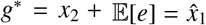, and the first equality of Eq. (8) results.

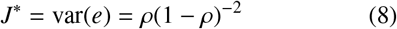

The second equality emerges by taking moments from the distribution of customers in an *M*|*M*|1 queue, which is 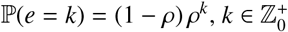 [27].

The distortion rate function, *d*, of a channel defines the minimum distortion achievable when transmitting information at a given rate [12]. It measures the optimal error properties of that channel. In [18] it was shown that for a Poisson channel, the optimal distortion at small *ρ* is *d* ≈ *ρ*. From Eq. (8) it is clear that lim_*p*→0_ *J*\* = *d*. This asymptotic equality defines the asymptotic optimality of birth-following, and means that, at the limit, it is as good as the best codec possible.

The relative performance of birth-following is evaluated by finding *ψ*\* = *j*\*/*D*. A *ψ*\* < 1 means that a better precision than the distortion bound has been achieved. Combining Eq. (7) and Eq. (8) gives Eq. (9).

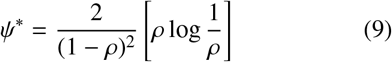

Observe that lim_*ρ*→0_ *ψ*\* = 0 because lim_*ρ*→0_ − *ρ* log *ρ* = 0. Thus, there exists a *ρ* = *a* such that *ψ*\* < 1. It can be shown that 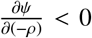, for all *ρ* ≤ *a* ≈ 0.199, so that *ψ*\* < 1 over this entire parameter regime. This contradicts the expected relation between *J*\* and *D* from Section 2.2, and was attributed to the diffusion approximation inherent in deriving *D* [17]. The sharp discreteness of birth-following allows it to operate beyond the limits of this approximation. This completes the proof of Theorem 1. This line of reasoning will underpin many of the estimation problems solved in this text.

Birth-following will be shown to quite generally outperform the bound because of the term −*ρ* log *ρ*. This term reveals an interesting connection between information and relative precision that validates *ψ*\* as a key metric. The queue distribution 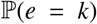 has entropy 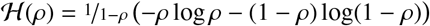 [12], which describes the most information that could be learnt from *e*. In the low *ρ* ≪ 1 regime 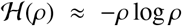 and 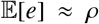. Combining these with a known inequality on −*ρ* log *ρ* from [12] yields Eq. (10).

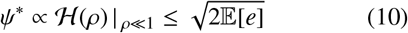

Eq. (10) states that relative precision is proportional to the maximum information in the queue and scales with the square root of the mean queue length, which can be thought of as a mean signalling threshold. Eq. (10) connects queueing, information and signalling. Note that all the expressions here depend on a single, dimensionless ratio, *ρ* = *u*/*f*_max_. All results are therefore scalable to any biological settings satisfying this ratio.

## 3. Results

### 3.1. Minimum Time Encoding

Birth-following is a heuristic encoder that was shown to be asymptotically optimal in [18] i.e. as *ρ* = *u*/*f*_max_ → 0 there is no better way of embedding information about *x*_1_ in *x*_2_. This result, however, provides no assessment of its biological importance, or if it this encoder is useful away from this theoretical limit. Here, by examining a class of stochastic control problems, birth-following is proven to be the fastest (minimum-time) strategy for signalling on a threshold, even outside the low *ρ* region. This establishes birth-following as a meaningful encoding paradigm, links it to biologically sensible timing problems [15] and motivates subsequent use of this strategy to assess the limits of pathway precision.

Consider the space of encoders, 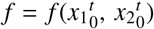, with full knowledge of the history of the target and signalling molecules. Assume that *f* only changes at molecular event times, and is constrained to *f*_min_ ≤ *f* ≤ *f*_max_ with 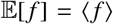. This makes biological sense as interevent decisions would require knowledge of an external clock and the next event time cannot be predicted exactly [29]. This assumption and the Markov nature of *x*_1_ and *x*_2_ transforms the search for an optimal *f* into a Markov policy design problem, over *e* = *x*_1_ − *x*_2_ [30].

As event times are unpredictable 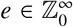 and *f*(0) = 0 = *f*_min_ (there is no point acting when *e* = 0). Designing *f*(*e*) is equivalent to controlling the random walk of *e* along its states, with the aim of forcing *e* to 0 as quickly as possible. This is a minimum time problem that is analogous to controlling the service rate of a queue to minimise congestion [29] [30]. All possible service rates are discretised within [0, *f*_max_]. The control law or policy *f* is a mapping from the queue length to these service rates. Optimal control theory can be used to find this mapping. In [29] an optimal policy, *f*\*, satisfying Eq. (11) for 0 ≤ *t* ≤ *τ* was derived.

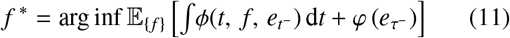

Here *t*^−^ means infinitesimally before time *t, ϕ* is a cost on the queue length and *φ* is a terminal cost. If *ϕ* = *af* + *e* i.e. it depends linearly on the queue length (*e*) and the service rate (*f*), and *φ* = 0, then the bang-bang controller of Eq. (12) emerges [29].

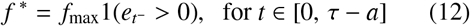

Bang-bang controllers are a class of policies that switch between extremes in response to the controlled variable (i.e. *e*). In this birth-process encoder design problem there is no service rate cost (*a* vanishes) and *J* is nondecreasing in *e*. Minimising *ϕ* thus achieves the mmse *J*\*, for a fixed *τ*, and Eq. (12) implies birth-following.

Further, consider the cost 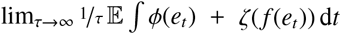 with *ϕ* and *ζ* as queue and service rate charges. If *ϕ*(*e*) is non-decreasing and convex in *e* with *ϕ*(0) = 0 and 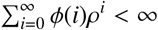 then the *f* that minimises this cost increases with *e*. This result from [31] and [30] is called a monotone optimal policy. The mse criterion satisfies these conditions with *ζ* = 0 for the encoder design problem. The lack of a service charge means *f*_max_ can be chosen when *e* = 1. Monotone optimality then requires that *f*_max_ also be chosen for *e* > 1. Hence *f*\* converges to birth-following. None of these results depend on *ρ*, beyond requiring a stable queue.

Interestingly, this monotone optimal argument also holds for complex networks of queues [32]. If the total queue cost is *ϕ* = ∑_*j*_ *ϕ_j_*, with *ϕ_j_* as the *j*^th^ queue cost, then a network of birth-following encoders is optimal [33]. As this network only consists of *M*|*M*|1 queues, it is a Jackson network [27]. Jackson networks are special because their constituent queues can be treated independently and in isolation. This observation, together with the fact that optimal results for Jackson networks also tend to hold for more arbitrary distributions [34], motivated the work in Section 3.2 and Section 3.3.

Thus, birth-following is a robust bang-bang rate controller. In [35] the problem of controlling the rate of a point process to maximise the probability of getting *N* events in time *τ* is studied. Using the cost in Eq. (11), [35] found that the minimum time policy is bang-bang over [0, *f*_max_]. Birth-following is therefore a minimum time encoder. Eq. (6) maximises the speed at which *x*_2_ can attain some *x*_1_ threshold, at any *ρ* = *u*/*f*_max_. As a result, birth-following could be important for realising fast and precise event timing in cellular transduction, especially when signals depend on thresholds [15]. These properties make birth-following ideal for probing whether or not capacity is a reliable proxy for defining precision on signalling pathways.

### 3.2. General Birth-Following

Having established the wider, meaningful optimality of birth-following, its behaviour relative to the capacity-based bounds of [1] is investigated. Previously, the *x*_1_ birth rate, *u*, was constant. This models constitutive expression in which a gene is continuously on, and commonly describes the behaviour of many housekeeping genes [36]. Here Theorem 1 type results are shown to hold for more complex and arbitrary *x*_1_ birth models, suggesting that in many instances existing bounds do not fully characterise achievable estimation precision. Two main 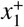 generalisations are developed.

First the birth rate of *X*_1_ is allowed to be any timevarying function, *λ*(*t*), constrained so that *u*/*ϵ* ≤ *λ* ≤ *u*, with *ϵ* > 1. Then *x*_1_ ~ Po(*λ*), and a Poisson channel with constraints 〈*f*〉 ≥ *u* and max(*f*) ≤ *f*_max_ is fixed. A utilisation *ρ* = *u*/*f*_max_ is also defined. This description encompasses a rich set of dynamical behaviours including (i) bursty models of gene expression that involve on-off switches (e.g. between *u*/*ϵ* and *u*) [36], (ii) external or environmental influences that change the constitutive transcription rate [37], and (iii) arbitrary positive or negative feedback controlled expression, in which *λ*(*t*) varies between its limits as a function of *x*_1_ [15].

To prove relative performance, hypothetical species at the limits of *λ* are needed. Let *Y*_0_ be one such species with constant birth rate *u* and population size *y*_0_. Then *x*_1_ can be constructed from *y*_0_ by independently accepting each 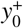 as a newly arriving customer in the 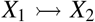 queue with probability *λ*(*t*)/*u* [38]. This queue is a consequence of *X*_2_ implementing birth-following. This means *y*_0_ ≥ *x*_1_. If *e*_0_ = *y*_0_ − *x*_2_ is the number of customers in a hypothetical 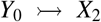 queue, then *e* = *x*_1_ − *x*_2_ ≤ *e*_0_. Computing the mmse on these queue lengths gives 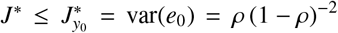. 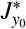 is from Eq. (8) as the 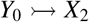 queue is *M*|*M*|1.

Denote another hypothetical species as *Y*_1_ with *y*_1_ ~ Po(*u*/*ϵ*). Its diffusion approximation is a scaled version of Eq. (5), and its bound, *D*_*y*_1__ = (−2*ϵ* log *ρ*)^−1^. If the unknown bound for *X*_1_ is *D* then *D* ≥ *D*_*y*1_ (*X*_1_ has a larger mean and variance than *Y*_1_, and is hence more difficult to estimate on the same fixed capacity channel). This also follows from known data processing inequalities for thinned Poisson processes [39]. Combining 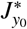 with *D*_*y*_1__ leads to an inequality on *ψ*\* for the 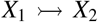 system, which is given in Eq. (13).

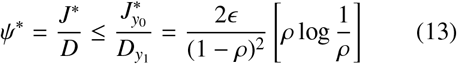

The −*ρ* log *ρ* factor means that *ψ*\* < 1 exists and so birth-following outperforms the capacity-based bound under finite, arbitrary, time-varying birth rates.

In the original 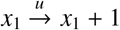 case, the inter-birth event (or waiting time) distribution is exponential with mean 1/*u*. The second generalisation investigated here relaxes this exponential waiting time to any distribution with the same mean. Births are still independent and identically distributed but *x*_1_ no longer conforms to a Poisson process description, and therefore no longer has to satisfy 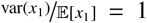. Non-exponential waiting distributions have been used to model burstiness and memory in transcription, as well as multi-step (e.g. gamma distributed) delays in protein production [40] [41].

Applying birth-following with *e* = *x*_1_ − *x*_2_ leads to a *G*|*M*|1 queue instead of the usual *M*|*M*|1. The ‘G’ indicates that the customer arrival distribution is generalised instead of exponential. The *G*|*M*|1 queue length distribution is 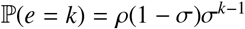 [27], with 0 ≤ *σ* < 1 derived from the chosen generalised arrival distribution. At *σ* = *ρ* the *M*|*M*|1 case is recovered. Since the mmse is unknown for this estimation problem, the mse is computed instead, resulting in Eq. (14).

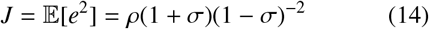

The channel constraints are unchanged so *D* is from Eq. (7). The gives *ψ*\* ≤ *ψ* = *J*/*D* as shown in Eq. (15).

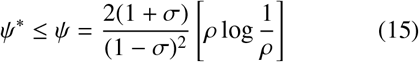

Enumerating over *σ* characterises the birth-following performance of all possible generalised distributions. Since every possible arrival distribution has *σ* < 1 [27], and the −*ρ* log *ρ* factor appears, then over some region of small *ρ, ψ*\* ≤ *ψ* < 1 is achieved as in Theorem 1. The capacity therefore provides a misleading assessment of achievable precision for a wide range of bimolecular estimation problems. The next section considers whether this observation holds for more complex networks.

### 3.3. Arbitrary Signalling Networks

The synthesis rate of the target molecule, *X*_1_, was previously generalised. Here the molecular network that is estimating *x*_1_ is generalised instead. This models complex pathways where several species are involved in signalling (e.g. molecular relays with scaffold proteins) or where multi-step delays are present. Similar, though more complex queue networks (e.g. with death events or feedback) have been used to study the lac operon and enzymatic correlations [42] [43]. While the approach here is simpler, the demonstrated performance of birth-following networks suggests its usability in synthetic circuit design (see Appendix B). Related results that include deaths are in Appendix A. Achievable precision is compared to that predicted by pathway capacity.

Consider arbitrarily configured molecular cascades with *n* − 1 hidden species, {*X*_2_,…, *X_n_*}, that estimate an upstream *X*_1_. Let only the final output of this network, *x*_*n*+1_, be observable. Each *X_j_* for 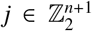 encodes its input using birth-following. An *M*|*M*|1 queue network results. As every encoder is independently constrained, each queue can have a different utilisation. Only feedforward networks are considered so that the signalling architecture is composed of splits and joins. The feedforward assumption is common in intracellular signalling analyses [15]. Burke’s theorem states that the output process of any *M*|*M*|1 queue is Poisson with rate equal to its input process [28]. Consequently, *x*_*n*+1_ must have rate *u*, by the principle of local balance [27].

A network with four intermediate species is given in Fig. 3 to illustrate the types of signalling architectures possible. Fig. 3 has five queues with parallel and serial connections. The *j*^th^ queue has length *e_j_* and utilisation *ρ_j_* < 1. The splitting process on the output of the *e*_1_ queue is a random Poisson thinning with probability *p*. The joining process that is the input of the *e*_5_ queue is a Poisson superposition [27]. All species have Po(*u*) statistics by Burke’s theorem. It is necessary to first derive capacity-based bound for these arbitrary networks.

**Figure 3:**
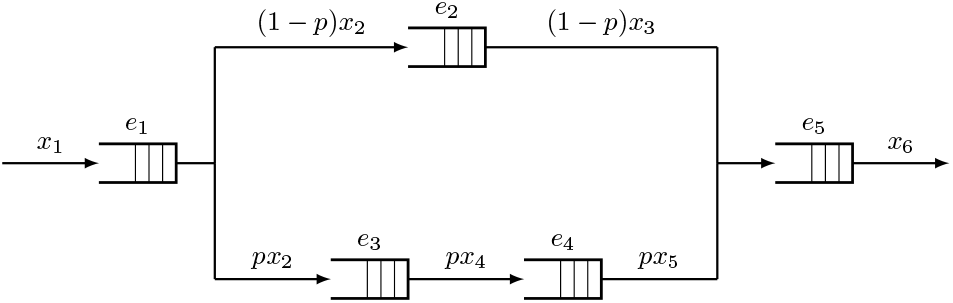
Example signalling network. Information about the target population, *x*_1_, is indirectly transferred to the observed population *x*_6_ by four intermediates. The network splits *x*_2_ (*p*-thinning) and then joins the branches as an input to the final queue (superposition). Birth-following is applied throughout so each queue is *M*|*M*|1.

Any such network can be modelled as a set of effectively serial links, each with rate *u*. Fig. 3 has three: *e*_1_, *e*_5_ and an effective one with the parallel queues. Every network always has at least one truly serial queue due to *x*_*n*+1_. While the capacity, *C_n_*, of an arbitrary cascade with *n* queues is unknown for *n* > 1, it is no greater than that of its most restrictive single, serial Poisson channel [44]. Without loss of generality it is assumed that the *e_n_* queue has the largest utilisation in the network, *ρ* = max_*j*_*ρ_j_*. Consequently, *C_n_* < *C*_1_ = −*u* log *ρ*, and the bound *D_n_* > *D*_1_ = (−2 log *ρ*)^−1^ (see Eq. (7)).

The *j*^th^ queue has stationary distribution 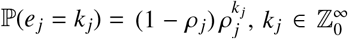. The effective error between the target and observed species, *e* = *x*_1_ − *x*_*n*+1_ is of interest. Solving across the network, it is found that in general 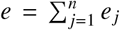. Jackson’s theorem states that the *e_j_* are mutually independent so 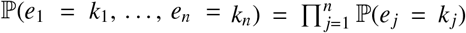 [27]. The optimal decoder is 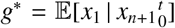 [25]. Let the mmse be 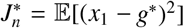. Then 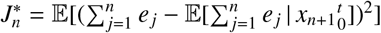. Using a corollary of Burke’s theorem, which states that queue *e_j_* is independent of its past departures [28], gives 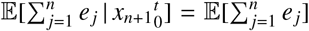. This implies that 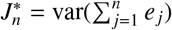, which when expanded gives Eq. (16).

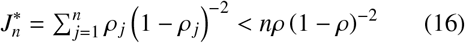

The optimal decoder is 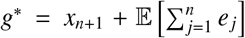. The network performance ratio, 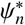, is derived as Eq. (17).

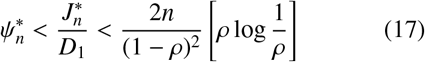

Intriguingly, 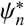 is bounded by a Theorem 1 type expression. Hence, for any network the bound of [1] will be violated by birth-following codecs at low maximum utilisation ρ. Consequently, over a large space of signalling networks, the capacity does not provide a complete description of the best achievable precision.

### 3.4. Poisson Sampling at Infinite Capacity

Birth-following broadly outperforms known capacity bounds when *ρ* is small. This corresponds to the maximum constraint (*f*_max_) being large relative to the mean (〈*f*〉) one. However, birth-following fixes 〈*f*〉 = *u* (*M*|*M*|1 property). Here this equality is relaxed, and the limit, *f*_max_ → ∞, of the conditions under which *J*\* < *D* is achieved is scrutinised. This results in unlimited information transfer (*C* → ∞) and perfect timing precision (signalling events can respond immediately to 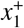). Both Poisson and Gaussian channel models predict perfect estimation mmse (i.e. *D* → 0) at this limit [13]. Comparing *J*\* to this prediction informs on how well *C* appraises precision in an information rich setting.

While transmitting unlimited information may seem impractical (real cells have finite rates), this analysis has several applications. First, it models the extreme of rapid cellular signalling, where *X*_2_ is much faster than *X*_1_. This situation is common in genetics and enzyme kinetics, where time scale separation methods are often used to eliminate the fast reactions from analysis [45]. This is equivalent to setting *f*_max_ → ∞. Second, infinite reaction rates may be explicitly assumed for simplicity. In [46] and [47] this is done to model fast binding between ribosomes and mRNA or substrates and catalysts.

Since signalling is without delay the estimation problem reduces to one of sampling and reconstructing the *x*_1_ ~ Po(*u*) population under the mean constraint 〈*f*〉 = *u*/*b*, with *b* > 0 as a mismatch parameter. Sampling is equivalent to encoding and reconstruction is the analogue of decoding. At any sampling time, *τ, x*_2_(*τ*) = *x*_1_(*τ*) and between samples *x*_2_ does not change. As a result *x*1 is referred to directly. For simplicity *x_t_* is used for *x*_1_(*t*) (the time index is more important here). Without loss of generality *x*_0_ = 0 is assumed. The aim is to design an optimal sampler or stopping strategy, *f*, that minimises mse while satisfying 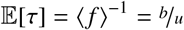.

The sampling time, *τ*, is a stopping time with respect to 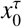, and *M_t_* = *x_t_* − *ut* obeys the optional sampling theorem [48]. This states that stopping times satisfy 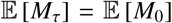 and gives 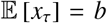 when combined with the 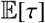 constraint. This means that only *f* functions that sample, on average, every *b* births of *X*_1_ are valid. The optional sampling theorem also applies to 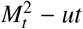 [48]. The further relation 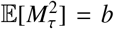 results. The conservation law of Eq. (18), which linearly trades between the variance of the samples and sample times, is derived by expanding this and noting that 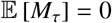.

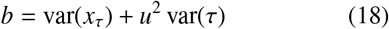

The mse of reconstruction for some decoder *g* is 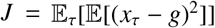 [49]. The *τ* subscript emphasises that averages are now across stopped trajectories. This is an ensemble equivalent to previous mse expressions. The optimal decoder 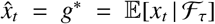 achieves mmse *J* = *J*\*, with 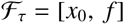 as the total causal information available (*t* ≤ *τ*), due to the Markov nature of *X*_1_ [24].

Before treating birth-following, deterministic protocols, which sample at a fixed *τ*, are examined. This is known as periodic sampling [50]. Let the encoder here be *f_d_*(‘*d*’ indicates deterministic). As *f_d_* has no information about *X*_1_ then 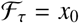 and 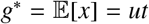. This gives 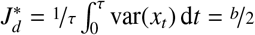. In this case *b* = *uτ* and var(*τ*) = 0 and so var(*x_τ_*) = *b* by Eq. (18). This scheme therefore maximises the sample variance.

A birth-following type protocol, called b-following, can be constructed by sampling every *b* events. This forces var(*x_τ_*) = 0 so that var(*τ*) = *b*/*u*^2^, is maximised (other extreme of Eq. (18)). This classes *b*-following as an adaptive sampler [50]. Adaptive samplers are known to improve upon deterministic schemes, such as the periodic one, by exploiting informative events [51]. At *b* ≤ 1 birth-following is recovered as an adaptive sampler. Biologically, *b*-following also makes sense since signalling on a threshold of molecular accumulation is a common motif in genetic networks [20] [15].

The optimal decoder for *b*-following over *t* ≤ *τ* is 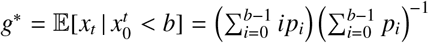 with 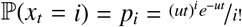. Solving gives Eq. (19).

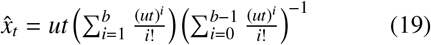

Eq. (19) satisfies 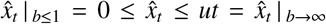. Interestingly, these limits recommend a transition in decoding from maintaining the last sampled value (here *x*_0_ = 0) when *b* ≤ 1 to a linear, uninformative function that matches the periodic sampler decoder as *b* → ∞. This exemplifies how signalling rate constraints can alter the definition of good performance.

If *t_i_*:= inf{*t: x_t_* ≥ *i*} then *t_i_* ~ Erlang(*i, u*) with density 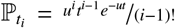 and (*t*_0_, *t_b_*) = (0, *τ*). The mmse is 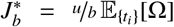 with expectations taken over 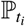 and 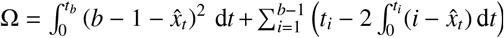. These expressions yield 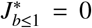. Since *D* = 0 here, capacity is a reliable proxy for precision when *b* ≤ 1. For 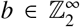 performance is not so promising. This is unsurprising as samples are taken less frequently than *x*_1_ births and the optimal reconstruction, 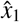, becomes dependent on (imperfectly) interpolating between these samples [52]. As a result 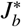 increases with *b*.

At large *b*, 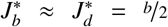 and the extra knowledge from adaptive sampling dissipates. At *b* = 2, 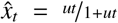 and Ω is 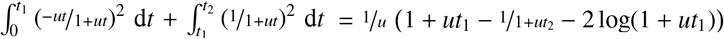. Using 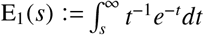 gives 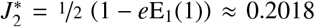. The *b*-following approach at *b* = 2 is illustrated in Fig. 4. Adaptive samplers at *b* > 1 therefore cannot achieve perfect precision, even with infinite capacity.

**Figure 4:**
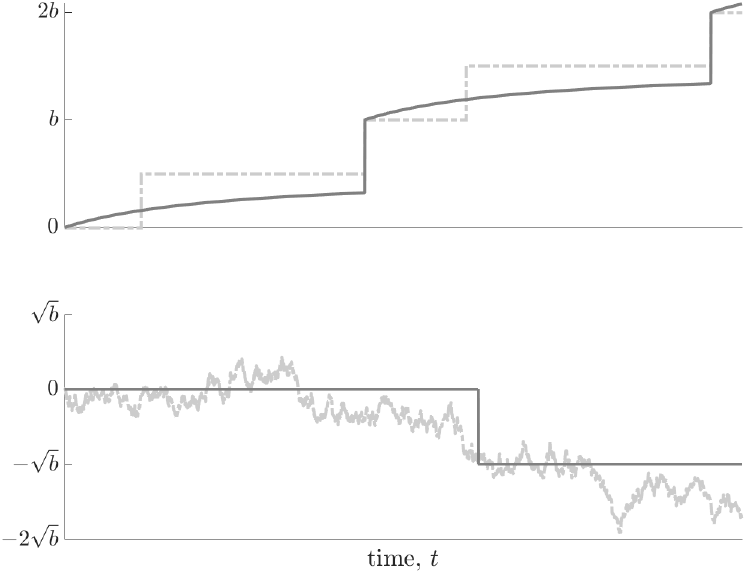
Fast codecs. The target molecule, *x*_1_ (dashed in both panels) is sampled and reconstructed (solid in both panels) over Poisson channels with infinite capacity. Here samples can be taken instantaneously (*f*_max_ → ∞) but the average sampling time is set to *b*/*u*, with *u* as the *x*_1_ synthesis rate. The top panel shows the optimal adaptive *b*-following codec with *b* = 2 for *x*_1_ as a birth-process. The bottom panel illustrates the optimal adaptive threshold sampler for any *b* with *u* = 1, but with *x*_1_ − *ut* as a standard Wiener process (see main text).

For comparison, a mmse of 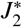 is achievable on a finite *f*_max_ birth-following channel with just *C* ≈ 0.4616 nats. Thus, higher capacities do not always guarantee improved precision. This contrasts the intuitive relationship between information and estimation found in both [1] and Gaussian channel descriptions. This reflects the fact that Gaussian capacities only depend on a single parameter (snr), while Poisson ones are sensitive to two independent constraints: 〈*f*〉 and *f*_max_. Appendix C shows that the many combinations of (〈*f*〉, *f*_max_) yielding the same *C* can lead to very different notions of absolute precision. Knowing the channel constraints is vital.

### 3.5. Removing the Diffusion Approximation

The inability of the capacity bound, *D*, to completely specify realisable precision has been proven. This has been largely attributed to the diffusion approximation of *X*_1_ inherent in *D* [17]. Here this approximation is made exact, and *f*_max_ → ∞ is considered as in Section 3.4. Since *C* → ∞, the estimation problem is now one of sampling and reconstruction. Let *x_t_* be a Brownian motion with drift that describes the target population at time *t*, with *x*_0_ = 0, so that 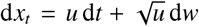 (Eq. (5)) holds. Designing samplers for *x_t_* under these conditions, at which *D* = *u*/2*C* → 0 [1], is investigated.

The goal is to optimally sample and reconstruct *x_t_* as a function of the mean encoding constraint 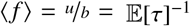, with *τ* as the sample time. Using the ergodic Markov nature of *x_t_*, the mmse can be computed as the ensemble average: 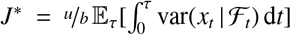 [49] [52]. Here F_t_ is all the causally observable information about *x_t_*. Sampling and reconstruction is equivalent to encoding and decoding in this problem. Periodic sampling, in which samples are taken every *τ* time units with var(*τ*) = 0 is first examined. As the sampler is independent of *x_t_*, 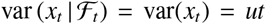 and so 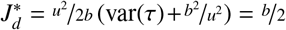, with ‘*d*’ for deterministic.

Adaptive or event based sampling can improve performance by capitalising on informative events in *x_t_* [50]. What signifies such an event, however, is not as clear as when *x_t_* ~ Po(*u*). Since the sampling problem with drift is no different from one without it [51], the associated process, *y_t_*, obeying 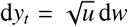, is examined. Further, *u* = 1 is set without loss of generality (time can be rescaled linearly to recover the 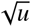). Such changes maintain *D* = 0 and make sampling results from [49] and [50] applicable. These state that it is mmse optimal to sample every time *y_t_* crosses a symmetrical threshold of ±*r*. This crossing is the informative event.

If *τ*_1_ is the first crossing time then 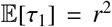 [48] and 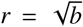. The optimal decoder under this sampler holds *ŷ_t_* at the last sample value before *t*. Fig. 4 shows *ŷ_t_* with a threshold sampling event leading to a stepwise shift in the reconstructed estimate (bottom panel). The mmse, 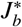, for this scheme is computed in Eq. (20), which is the limit of a relation derived in [49] using optimal stopping theory, with *e* as the natural exponent.

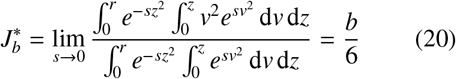

In [50] this threshold based sampler was shown to imply a stationary conditional variance of *r*^2^/6. As a result of this and Eq. (20), 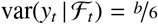, for any time scaling. Since the difference between *x_t_* and *y_t_* is simply a drift term then Eq. (20) gives the mmse for *x_t_* as well.

At finite 〈*f*〉 (*b* > 0) it is thus impossible to reconstruct *x_t_* perfectly, in contrast to *D*. While the optimal threshold sampler is not analogous to birth-following, it is similar to a toggle based encoder in [17], which outperforms the bound for birth-death processes. The persistence of these mismatches, even when the diffusion approximation is no longer a source of error, highlights the difficulty in characterising the precision limits of birth-processes with capacity. The relationship between information and achievable precision thus remains complicated, and potentially misleading.

### 3.6. Example Applications

Birth-following is an optimal strategy that has been found to broadly disrupt the known limits of achievable precision. Examples of real or modelled biosystems, to which these results can provide practical insight follow.

(a) Proportional synthesis. In bacteria and eukaryotes the subunits of many multiprotein complexes are produced in stoichiometric quantities [53] [54]. This sharply constrained production, which does not employ feedback, is called proportional synthesis and relies on finely tuned birth rates that are proportional to the desired subunit quantities. Tight coupling is necessary because synthesis is costly and subunit population mismatches can lead to misfolding or aggregation errors [53]. This stoichiometric requirement means that complexes with subunits in a 1:1 ratio would have equal subunit (mean) synthesis rates. This motif applies to stable proteins, so death rates can be neglected [53].

The mechanism underlying proportional synthesis remains unknown [54]. If a 1:1 complex with proteins *X*_1_ and *X*_2_ (a heterodimer) is considered, then a minimum model for efficiently generating stoichiometric quantities could be: *x*_1_ ~ Po(*u*(*t*)), *x*_2_ ~ Po(*f* (*t*)) with 〈*u*〉 = 〈*f*〉. This model, which focuses on the subunit coupling, abstracts multistep processes that could lead to the formation of individual proteins by making the synthesis rates arbitrary and time varying. Such an abstraction was found sensible in [55] (see Appendix B).

The *X*_1_ and *X*_2_ birth (synthesis) rates must be coupled else the mmse between their populations could explode. Under these conditions birth-following is the optimal coupling solution (see Section 3.2), minimising mismatch, while maintaining the mean equality constraint. This would support the belief that proportional synthesis minimises the negative effects of uncomplexed subunits while maximising synthesis efficiency [53] and could help characterise the limiting precision of this motif. Other stoichiometric ratios of 1:b could be realised by modifying birth-following so that it signals *b* times for every *x*_1_ event, as explored in [18].

(b) Targeted protein degradation. ClpXP is a protease in E. coli that catalytically degrades mistranslated proteins. [47]. It possesses multiple sites, to which substrate proteins will often simultaneously bind. ClpXP achieves efficient degradation by processing substrate molecules in turn, creating a waiting line over its sites. This has been naturally modelled as a queue with the protein molecules as customers, and ClpXP as the server, which processes (i.e. degrades) the customers in sequence [56] [47]. If *e* is the number of queueing customers then the model used in [47] is 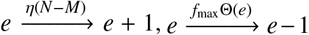, with *N* and *M* counting the total number of binding and unoccupied sites, *η* as a rate constant and Θ(*e*) as a step function that is 1 when *e* > 0.

This conforms to the framework in Section 3.2 with *u* = *ηN* and effectively realises a birth-following through Θ(*e*). However, this only holds if *f*_max_ is sufficiently large so that *e* ≤ *N*. When this model is extended to allow for multiple protein classes then a more complex queue network is obtained to which results from Section 3.3 and Appendix A apply. The properties of birth-following explain several observations such as the low latency of the waiting line, the approximately exponential departure times of substrate from ClpXP, even in cases involving complex networks and the high temporal precision of the response, which targets aberrant proteins for fast degradation but then immediately turns off once no more excess substrate exists [56] [47].

Birth-following could provide a useful reference for assessing the mmse achievable by these targeted degradation systems, where *X*_1_ is the aberrant protein and the *X*_2_ is the degraded substrate. Minimising the mse corresponds to removing the aberrant protein with minimum delay. The usefulness of birth-following as an explanatory mechanism may hold more generally as many other enzymes may exhibit queue like processing [56].

Many further applications exist. Models describing the observational limits of fluorescent proteins [36], accurate protein production from a single transcript [41], coding performance in sensory transduction [3] [14], and rate precision in phylodynamic processes [57] [58] are all amenable to this analysis. This follows from the ubiquitous importance of discreteness, stochastic delay and timing accuracy in biosystems. Birth-following could serve as a null hypothesis here for contextualising observed performance against realisable limits, providing insight without the need for comprehensive biological knowledge. Systems not implementing birth-following under the relevant constraints, for example, cannot be optimising for speed. The capacity would not be a reliable proxy for precision in these systems.

## 4. Discussion

Noise plays a formative role in cell biology, often shaping the mechanisms and motifs that regulatory molecular networks have to implement in order to attain precise functionality [4]. Many of these mechanisms, such as those behind stoichiometric synthesis or coupled feedback, are still unknown or under investigation [59] [22] [60]. Information theory has proven useful in this context by allowing initial analyses of what signalling pathways can and cannot achieve without requiring detailed knowledge of the complex molecular network involved [3] [8]. Specifically, studies have computed the capacities of signalling pathways and used this to support or refute hypothetical motifs about the achievable precision on that pathway [7] [61] [5].

While several important insights have emerged from this approach, the relationship between capacity and precision is still unclear and sometimes deceptive, especially when the discreteness and causality of the information transmitted on a pathway are taken into account [1] [14] [17]. This work cautions against overinterpreting this relationship by clarifying, for a class of simple but non-trivial birth-process networks, what is an optimal signalling mechanism (encoder) and how it redefines the plausibly achievable estimation precision on a signalling pathway of known capacity.

By taking a discrete, Poisson channel approach to signalling, it was found that birth-following, a scheme previously thought heuristic [17], is actually the unique minimum time encoding solution for birth-process estimation (Section 3.1). Birth-following behaves like a molecular switch or relay [62], activating at maximum speed on an observed target birth, and then deactivating with minimum delay once no events remain to be signalled. This scheme achieves the most precise signal timing possible (within its rate constraints), and could be an important null hypothesis for evaluating observed biomolecular motifs. A regulatory birth-process network that fails to even approximate birth-following likely does not prioritise response speed.

The optimal properties of birth-following also make it an excellent candidate for synthetic circuit design, especially when multiple, precisely timed signals are of interest [62]. This encoder could potentially be realised either exactly using a queueing approach or approximately using circuits that implement sharp, sigmoidal Hill functions (see Appendix B) [56]. Interestingly, birth-following, which can also be thought of as a coupled random telegraph, would not have been the optimal encoding choice had a Gaussian channel model been used. While random telegraph signals achieve the capacity of Poisson channels [26], only normally distributed inputs can maximise information flow on Gaussian ones [12]. This hints at how continuity approximations can lead to misleading conclusions [14].

Given its optimality, birth-following was used to probe how realisable precision compares to the best known theoretical bounds from [1], which quantify how Poisson channel capacity specifies maximum precision. Birth-following was found to achieve higher precisions (lower mse values), provided the dynamic range of the channel, *f*_max_/〈*f*〉, was large enough and the mean signalling rate 〈*f*〉 was no smaller than the mean target birth-rate. This violation persisted over wide generalisations to both the target molecule rate and the signalling network architecture (Section 3.2 and Section 3.3). Mismatches were maintained even when death reactions, which model molecular turnover, were included, though an extra channel was needed to compensate for the extra death noise (see Appendix A). These results greatly extend and contextualise those in [18].

Several biological systems, including those implementing proportional synthesis and targeted degradation (Section 3.6), lie within this generalised birth-following framework [56] [20] [53]. It appears that channel capacity is not sufficient to completely characterise, and in fact may underestimate, the best possible performance on a signalling pathway. This underestimation emerges from existing bounds using a diffusion process to approximate the dynamics of the target molecule (see Section 2.2). When *f*_max_/〈*f*〉 is large and 〈*f*〉 is no smaller than the average target birth rate, signalling events can respond to individual jumps in the target species. Target discreteness therefore cannot be ignored, and may be doubly important in cellular systems given that many molecular species are present in low numbers [4].

The 〈*f*〉 and *f*_max_ conditions that promote mismatches correspond to Poisson channels with high capacity. This suggests that the common belief of increased capacity implying increased precision still holds, just not always as derived in [1]. To challenge this assertion *f*_max_ and hence capacity were allowed to become un-bounded, resulting in the estimation problem reducing to one of sampling and reconstruction. In this case both the bound from [1] and standard Gaussian channel analyses predict that zero reconstruction error is possible.

By generalising birth-following to *b*-following and deriving optimal reconstruction functions, it was proven that perfect precision is impossible for any 〈*f*〉 smaller than the target birth rate (Section 3.4). To explore this observation further the sampling problem was reexamined with the target species conforming to a diffusion process so that the bounds of [1] become exact. By applying the known optimal threshold-based sampler for this problem [51], it was found that perfect precision can never be achieved at any finite 〈*f*〉 (Section 3.5). Channel capacity, *C*, therefore cannot be used as a bijective predictor of biological precision.

Analyses that compute *C* directly (e.g. by calculating the maximum mutual information of a pathway [3]) could potentially be misleading since the various combinations of 〈*f*〉 and *f*_max_ that map to the same computed *C* may impose very different restrictions on the attainable precision. For example, sometimes the same *C* and *f*_max_ correspond to both a 〈*f*〉 > *u* and 〈*f*〉 < *u* setting (see Appendix C). The bound of [1] would then overestimate the achievable precision in the first case, and underestimate it in the second. To truly gauge the precision achievable on a pathway, it is therefore more important to establish what signalling rate constraints hold. This aligns with the implication in [22] that more experimental investigation must focus on ‘how biochemical properties place constraints on individual motifs’.

## Acknowledgements

This work was partially funded by the Gates Cambridge Trust and the Control Group at the Department of Engineering, University of Cambridge. Joint centre funding from the UK Medical Research Council and Department for International Development under grant reference MR/R015600/1 is also acknowledged. The author thanks Glenn Vinnicombe and Paula Dobrinić for their thoughtful inputs and comments on this project.

## Appendix A. Death Reaction Networks

The pure birth-processes analysed in the main text are good models for the accumulation of stable molecules. Here a standard protein production-degradation model [2], 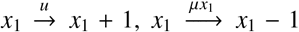, is used to examine how deaths alter the information structure and hence the form of optimal encoders. These reactions suggest that *x*_1_ can be thought of as the number of customers in an *M*|*M*|∞ queue with utilisation 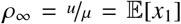 [27]. The ‘∞’ refers to an unlimited number of servers. The aim is to compute and compare *J*\* and *D* for this system.

The network of Fig. A1 uses two channels, employing birth-following between 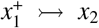 and 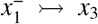. The combined scheme is termed birth-death following (*x*_3_ tracks molecular deaths). Here *x*_2_ and *x*_3_ are birthprocesses, and the *e*_1_ and *e*_2_ queues are both *M*|*M*| 1 by birth-following. Identical signalling rate constraints are placed on both channels. By applying Burke’s theorem to the *M*|*M*|∞ queue, it is found that customers enter both *M*|*M*|1 queues with rate *u* (see Section 3.3). The utilisation of both *e*_1_ and *e*_2_ queues is then *ρ* = *u*/*f*_max_.

**Figure A1:**
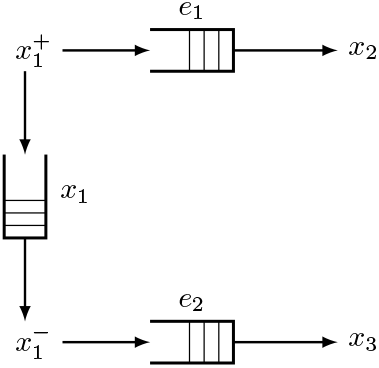
Birth-death following network. Signalling molecules *x*_2_ and *x*_3_ observe the target population births, 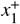, and deaths, 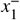. The *x*_1_ queue is *M*|*M*|∞ with number of customers equal to the target population size. The *e*_1_ and *e*_2_ queues are *M*|*M*|1 by birth-following.

Since two parallel channels are used the Poisson capacity doubles [1]. Using the queueing structure in Fig. A1 yields the new capacity bound of Eq. (A.1).

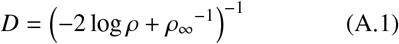

Let 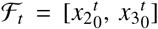 be all the observable information. The optimal decoder is 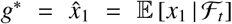. No extra information about one *M*|*M*|1 queue, given its output, is obtained by observing the output of the other *M*|*M*|1 [18] so 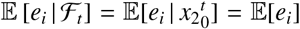 for 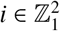. The last equality is from Theorem 1. The mmse is 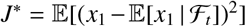 with 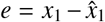. Expanding gives 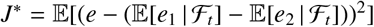.

Using previous equalities and that 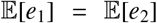 leads to 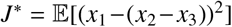. The optimal decoder is then the simple population difference 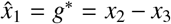. Substituting *e*_1_ − *e*_2_ = *x*_1_−(*x*_2_-*x*_3_), with *ω* = corr(*e*_1_, *e*_2_) and var(*e*_1_) = var(*e*_2_) = *ρ*(1 − *ρ*)^−2^ gives Eq. (A.2).

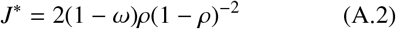

The correlation derives from the partial synchronisation of the *M*|*M*| 1 inputs and results in what is known as a Flatto-Hahn-Wright description [63]. The relative performance ratio, *ψ*\* = *j*\*/*D* then follows as Eq. (A.3).

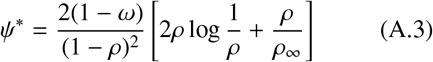

Since synchrony is never perfect: 0 ≤ *ω* < 1. Further, 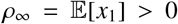. For small 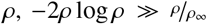 and 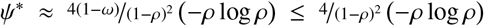. This generalises Theorem 1. Birth-death following therefore also outperforms the capacity bound, and is analogous to birth-following for a pure birth-process. However, its efficiency, 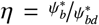 (‘*b*’ and ‘*d*’ signify births and deaths), is reduced due to the death noise, which increases *ψ*\*. Here *η* = 1/2 (1 – *ω*)^−1^ ≥ ‘/2, so up to 50% of the birth-following precision is lost due to deaths.

Deaths therefore change the information structure of the signalling problem, requiring two channel encoding. Single channel solutions are possible, but require complex, non-biological codecs [17]. Importantly, it is consistently found that birth-following is a high performing strategy and that capacity does not fully depict achievable molecular precision. Note that, practically, both births and deaths can be monitored by tracking molecular population numbers across time [64].

## Appendix B. Biological Realisability

Birth-following is an optimal encoder for problems in which a target molecule *X*_1_ must be sensed, monitored or tracked by a signalling one, *X_j_*. Mechanistically, it acts as a minimum time molecular switch or relay; activating on an event (*X*_1_ births), completing a signalling objective (effecting an *X*_2_ birth) and then deactivating with maximum speed. This allows it to achieve sharp timing (see Fig. 2), attain desired signalling thresholds quickly and make multiple signals efficiently by rapidly turning on and off [62]. Consequently, as it could be useful for synthetic genetic circuit design, the biological realisability of birth-following is examined.

Many biosystems (e.g. cooperative ligand binding or transcriptional regulation) implement non-linear, sigmoidal responses to target molecules that are described by Hill functions [65]. Eq. (B.1) defines a rising Hill function with maximum *f*_max_, input *e*, coefficient *h*(which controls its shape) and midpoint at *e* = 1.

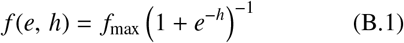

Fig. B1 shows that as *h* increases Eq. (B.1) converges to Eq. (6). Birth-following can therefore be easily approximated as a Hill function response to the excess in signalling over target molecules (i.e. *e* = *x*_1_ − *x*_2_).

Hill functions have previously been realised, but at moderate *h*. Toggle switch circuits can, however, model severe Hill type responses and so synthetically mimic Eq. (6) [19]. Birth-following can also be implemented using its queuing interpretation. While this has not been explicitly done yet, a similar queueing circuit has been designed, for ClpXP modelling (Section 3.6) in [56].

**Figure B1:**
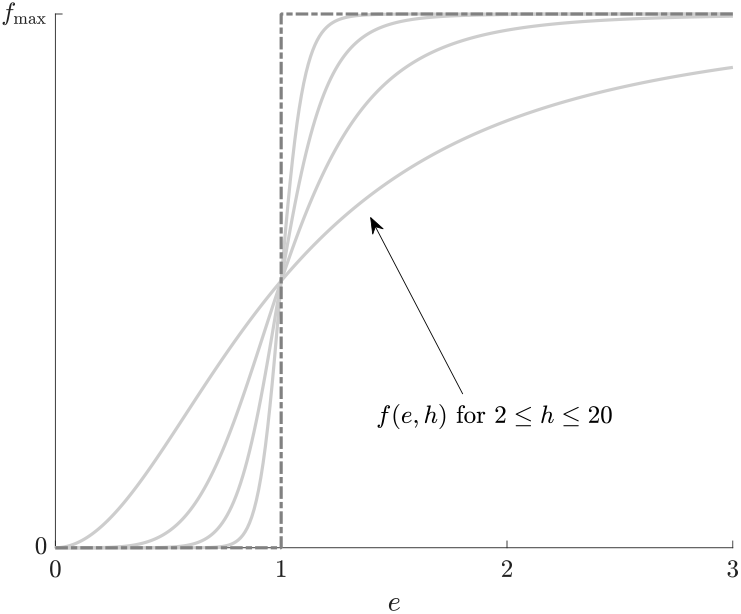
Hill function encoding. For a molecular count, *e*, Hill function encoders, *f* (*e, h*) are shown (solid grey) for a threshold value of 1, and with Hill coefficient, *h*. As *h* increases the Hill functions approximate a birth-following switch (Eq. (6)) (dashed grey).

One issue that synthetic circuits face is that of context dependence. While circuits are often designed to obtain some desired functionality, they may fail to do so once integrated into cellular processes due to changing environments, fluctuating in-cell conditions or unexpected coupling. These factors form the cell context [66]. An advantage of birth-following is that it maintains its properties (at least under the condition of fast signalling or low *ρ*) despite environmental fluctuations.

Consider a synthetic problem where birth-following is designed to allow *X*_2_ to monitor *X*_1_ under the belief that *x*_1_ ~ Po(*λ*). Let *z*, a stochastic environmental variable, unexpectedly modulate *X*_1_ so that in the cell Po(*λr*(*z*)) is the actual system, with *r*(*z*) as some function. This environmental effect can be marginalised so 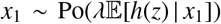 [55]. If this rate, which is now deterministic in *x*_1_ [25], can be bounded by a constant *u* then results from Section 3.2 apply. Birth-following is therefore context independent under bounded perturbations, and suitable for synthetic realisation.

## Appendix C. Mapping capacity to constraints

Experimental approaches to ascertaining the absolute precision of a signalling pathway often compute capacity, *C*, directly from empirical probability distributions [3] [7]. These distributions are used to calculate the maximum mutual information, which estimates the capacity. This is then mapped to a measure of precision (e.g. mse) by assuming either a Gaussian or Poisson channel model. Under both models a bijective relationship between capacity and precision exists [13] [1]. The latter model and its precision relationship (the distortion bound of [1]) represents the current state of the art, and has yielded several important insights.

However, as shown in Section 3.4 and Section 3.5 in the case of the Poisson capacity becoming very large, varying absolute precisions can be attained at the same capacity. Depending on the existing combination of mean, 〈*f*〉, and maximum, *f*_max_, signalling constraints, the bounds of [1] can either be violated or found valid. This work posits that this is a consequence of the non-bijective relationship between *C* and its constraints.

**Figure C1:**
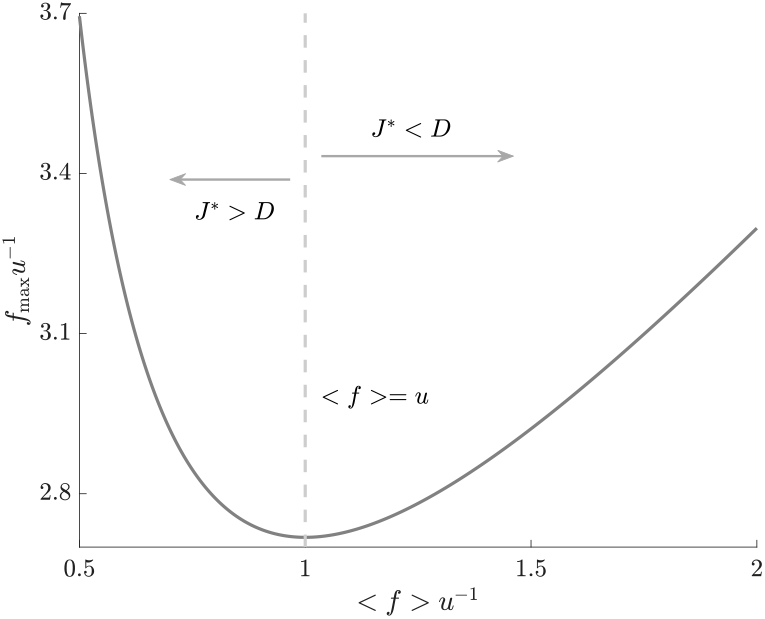
Non-bijective capacity-constraint relationships. All possible combinations of the mean (〈*f*〉) and maximum (*f*_max_) signalling rate satisfying the ratio *C/u* = 1 are shown. Knowing only the capacity (*C*) is insufficient for characterising absolute precision. Even specifying *C* and *f*_max_ is not enough as two different 〈*f*〉 values correspond to these settings. If *C* is very large (see Section 3.4), then these settings can lead to strikingly distinct beliefs about precision (demarcated by arrows with J* as the mmse and *D* the bound [1]).

For a given *C* with 〈*f*〉 = *ku* and *f*_max_ = *μu* the expression in Eq. (2) can be manipulated to obtain *μ* = *k e ^C/ku^*, with *e* as the natural exponent. Hence a range of signalling constraints can be combined to obtain the same *C*. This curve is given in Fig. C1 at *C/u* = 1. While *C* alone is clearly insufficient for characterising precision, the additional knowledge of *f*_max_ is still inadequate as two 〈*f*〉 values can be valid, one at *k* < 1 and one at *k* > 1. If *C* is very large then these conditions can correspond to the best achievable precision or mmse, *J*\*, either being over or underestimated by the bounds, *D* from [1]. This was shown in Section 3.4 with *b* = 1/*k*.

## Bibliography

[1] I. Lestas, G. Vinnicombe, J. Paulsson, Fundamental Limits on the Supression of Molecular Fluctuations, Nature 467 (2010) 174–8.

[2] D. Wilkinson, Stochastic Modelling for Quantitative Description of Heterogeneous Biological Systems, Nature Reviews: Genetics 10 (128).

[3] A. Rhee, R. Cheong, A. Levchenko, The Application of Information Theory to Biochemical Signaling Systems, Physical Biology 9 (2012) 045011.

[4] A. Eldar, M. Elowitz, Functional Roles for Noise in Genetic Circuits, Nature 467.

[5] C. Waltermann, E. Klipp, Information Theory Based Approaches to Cellular Signaling, Biochimica et Biophsica Acta 1810(2011) 924–32.

[6] J. Selimkhanov, B. Taylor, J. Yao, et al., Accurate Information Transmission through Dynamic Biochemical Signaling Networks, Systems Biology 346 (6215).

[7] R. Cheong, A. Rhee, C. Wang, et al., Information Transduction Capacity of Noisy Biochemical Signaling Networks, Science 334 (6054) (2011) 354–58.

[8] A. Hilfinger, T. Norman, G. Vinnicombe, et al., Constraints on Fluctuations in Sparsely Characterized Biological Systems, Physical Review Letters 116 (058101).

[9] M. Brennan, R. Cheong, A. Levchenko, How Information Theory Handles Cell Signaling and Uncertainty, Science 338 (6105) (2012) 334–335.

[10] S. Uda, S. Kuroda, Analysis of Cellular Signal Transduction from an Information Theoretic Approach, Seminars in Cell and Developmental Biology 51 (2016) 24–31.

[11] Z. Mousavian, J. Diaz, A. Masoudi-Najad, Information Theory in Systems Biology. Part II: Protein-Protein Interaction and Signaling Networks, Seminars in Cell and Developmental Biology (2016) 14–23.

[12] T. Cover, J. Thomas, Elements of Information Theory Second Edition, John Wiley and Sons, 2006.

[13] D. Guo, D. Shamai, S. Verdu, Mutual Information and Minimum Mean-Square Error in Gaussian Channels, IEEE Transactions on Information Theory 51 (4) (2005) 1261–82.

[14] K. Parag, G. Vinnicombe, Point Process Analysis of Noise in Early Invertebrate Vision, PLoS Computational Biology 13 (10) (2017) e1005687.

[15] K. Ghusinga, J. Dennehy, A. Singh, First-passage Time Approach to Controlling Noise in the Timing of Intracellular Events, Proceedings of the National Academy of Science 114 (4) (2017) 693–8.

[16] Y. Kabanov, The Capacity of a Channel of the Poisson Type, Theory of Probability and its Applications 26 (1978) 143–147.

[17] K. Parag, G. Vinnicombe, Single Event Molecular Signalling for Estimation and Control, European Control Conference (2013) 4166–71.

[18] K. Parag, G. Vinnicombe, Event Triggered Signalling Codecs for Molecular Estimation, 52nd IEEE Conference on Decision and Control (2013) 256–61.

[19] J. Teo, S. Woo, R. Sarpeshkar, Synthetic Biology: A Unifying View and Review Using Analog Circuits, IEEE Transactions on Biomedical Circuits and Systems 9 (4) (2015) 453–74.

[20] S. Gupta, J. Varennes, H. Korswagen, et al., Temporal Precision of Regulated Gene Eexpression, PLoS Computational Biology 14 (6) (2018) e1006201.

[21] M. Frey, Capacity of the Lp Norm Constrained Poisson Channel, IEEE Transactions on Information Theory 38 (2) (1992) 445–50.

[22] R. Gao, A. Stock, Overcoming the Cost of Positive Autoregulation by Accelerating the Response with a Coupled Negative Feedback, Cell Reports 24 (2018) 3061–71.

[23] D. Guo, S. Shamai, S. Verdu, Mutual Information and Conditional Mean Estimation in Poisson Channels, IEEE Transactions on Information Theory 54 (5) (2008) 1837–49.

[24] D. Snyder, Filtering and Detection for Doubly Stochastic Poisson Processes, IEEE Transactions on Information Theory 18 (1972) 91–102.

[25] D. Snyder, M. Miller, Random Point Processes in Time and Space, 2nd Edition, Springer-Verlag, 1991.

[26] M. Davis, Capacity and Cutoff Rates for Poisson Type Channels, IEEE Transactions on Information Theory 26 (1980) 710–715.

[27] L. Kleinrock, Queueing Systems Volume I Theory, John Wiley and Sons, 1975.

[28] P. Burke, The Output of a Queueing System, Journal of Operational Research Society 4 (1956) 699–704.

[29] R. Boel, P. Varaiya, Optimal Control of Jump Processes, SIAM Journal of Control and Optimization 15 (1) (1977) 92–119.

[30] R. Serfozo, Optimal Control of Random Walks, Birth and Death Processes, and Queues, Advances in Applied Probability 13 (1981) 61–83.

[31] T. Crabill, Optimal Control of a Service Facility with Variable Exponential Service Times and Constant Arrival Rate, Management Science 18 (9) (1972) 560–6.

[32] R. Weber, S. Stidham, Optimal Control of Service Rates in Networks of Queues, Advances in Applied Probability 19 (1987) 202–18.

[33] Z. Rosberg, P. Varaiya, J. Walrand, Optimal Control of Service in Tandem Queues, IEEE Transactions on Automatic Control 27 (3) (1982) 600–10.

[34] S. Stidham, Analysis, Design and Control of Queueing Systems, Operations Research 50 (1) (2002) 197–216.

[35] P. Bremaud, Bang-bang Controls of Point Processes, Advances in Applied Probability 8 (1976) 385–94.

[36] T. Lionnet, R. Singer, Transcription goes Digital, EMBO Reports 13 (4) (2012) 313–21.

[37] S. Nandi, A. Ghosh, Transcriptional Dynamics with Timedependent Reaction Rates, Physical Biology 12 (2015) 016015.

[38] P. Lewis, G. Shedler, Simulation of Non-homogeneous Poisson Processes by Thinning, Naval Res. Logistics Quart. 26 (3) (1989) 403–13.

[39] L. Wang, A Strong Data Processing Inequality for Thinning Poisson Processes and Some Applications, in: 2017 IEEE International Symposium on Information Theory (ISIT), 2017, pp. 3180–4.

[40] T. Jia, R. Kulkarni, Intrinsic Noise in Stochastic Models of Gene Expression with Molecular Memory and Bursting, Physical Review Letters 106 (2011) 058102.

[41] K. Josic, J. Lopez, W. Ott, et al., Stochastic Delay Accelerates Signaling in Gene Networks, PLoS Computational Biology 7 (11) (2011) e1002264.

[42] A. Arazi, E. Ben-Jacob, U. Yechiali, Bridging Genetic Networks and Queueing Theory, Physica A 332 (2004) 585–616.

[43] L. Tsimring, Noise in Biology, Reports on Progress in Physics 77 (2014) 026601.

[44] R. Sundaresan, S. Verdu, Capacity of Queues via Point Process Channels, IEEE Transactions on Information Theory 52 (6) (2006) 2697–709.

[45] J. Gunawardena, Time-scale Separation – Michaelis and Menten’s Old Idea, still Bearing Fruit, FEBS Journal 281 (2014) 473–88.

[46] W. Mather, J. Hasty, L. Tsimring, et al., Translational cross talk in gene networks, Biophysical Journal 104 (2013) 2564–72.

[47] P. Hochendoner, C. Ogle, W. Mather, A Queueing Approach to Multi-site Enzyme Kinetics, Interface Focus 4 (2014) 20130077.

[48] R. Serfozo, Basics of Applied Stochastic Processes, Springer Science and Business Media, 2009.

[49] M. Rabi, Packet Based Inference and Control, Ph.D. thesis, University of Maryland (2006).

[50] K. Astrom, B. Bernhardsson, Comparison of Periodic and Event Based Sampling for First Order Stochastic Systems, in: Proceedings of the 14th IFAC World Congress, 1999, pp. 5006–11.

[51] M. Rabi, G. Moustakides, J. Baras, Adaptive Sampling for Linear State Estimation, SIAM Journal of Control and Optimization 50 (2) (2012) 672–702.

[52] K. Parag, G. Vinnicombe, Constrained Adaptive Sampling and Causal Estimation of Stochastic Processes, Tech. rep., University of Cambridge (2015).

[53] G. Li, D. Burkhardt, C. Gross, et al., Quantifying Absolute Protein Synthesis Rates Reveals Principles Underlying Allocation of Cellular Resources, Cell 157 (2014) 624–35.

[54] J. Taggart, G. Li, Production of Protein-Complex Components is Stoichiometric and Lacks General Feedback Regulation in Eukaryotes, Cell Systems 7 (2018) 580–9.

[55] C. Zechner, H. Koeppl, Uncoupled Analysis of Stochastic Reaction Networks in Fluctuating Environments, PLoS Computational Biology 10 (12) (2014) e1003942.

[56] N. Cookson, W. Mather, T. Danino, et al., Queueing Up for Enzymatic Processing: Correlated Signaling through Coupled Degradation, Molecular Systems Biology 561 (2011) 1–9.

[57] K. Parag, O. Pybus, Optimal Point Process Filtering and Estimation of the Coalescent Process, Journal of Theoretical Biology (2017) 153–67.

[58] K. Parag, O. Pybus, Exact Bayesian Inference for Phylogenetic Birth-death Models, Bioinformatics 34 (21) (2018) 3638–45.

[59] C. Vogel, E. Marcotte, Insights into the Regulation of Protein Abundance from Proteomic and Transcriptomic Analyses, Nature Reviews: Genetics 13 (2012) 227–32.

[60] A. Levin-Karp, U. Barenholz, T. Bareia, et al., Quantifying Translational Coupling in E. coli Synthetic Operons Using RBS Modulation and Fluorescent Reporters, ACS Synthetic Biology 2 (2013) 327–36.

[61] I. Mian, C. Rose, Communication Theory and Multicellular Biology, Integrative Biology 3 (2011) 350–67.

[62] B. Alberts, A. Johnson, J. Lewis, et al., Molecular Biology of the Cell, sixth Edition, Garland Science, 2015.

[63] L. Flatto, S. Hahn, Two Parallel Queues Created by Arrivals with Two Demands, SIAM Journal of Applied Mathematics 44 (5) (1984) 1041–53.

[64] J. Yu, J. Xiao, X. Ren, et al., Probing Gene Expression in Live Cells, One Protein Molecule at a Time, Science 311 (5767) (2006) 1600–3.

[65] M. Santillan, On the Use of the Hill Functions in Mathematical Models of Gene Regulatory Networks, Mathematical Modelling of Natural Phenomena 3 (2) (2008) 85–97.

[66] S. Cardinale, A. Arkin, Contextualizing Context for Synthetic Biology – Identifying Causes of Failure of Synthetic Biological Systems, Biotechnology Journal 7 (2012) 856–66.

